# Tribus: Semi-automated discovery of cell identities and phenotypes from multiplexed imaging and proteomic data

**DOI:** 10.1101/2024.03.13.584767

**Authors:** Ziqi Kang, Angela Szabo, Teodora Farago, Fernando Perez-Villatoro, Ada Junquera, Saundarya Shah, Inga-Maria Launonen, Ella Anttila, Julia Casado, Kevin Elias, Anni Virtanen, Ulla-Maija Haltia, Anniina Färkkilä

**Author notes:** Correspondence should be addressed to A.F.

## Abstract

**Motivation:** Multiplexed imaging and single-cell analysis are increasingly applied to investigate the tissue spatial ecosystems in cancer and other complex diseases. Accurate single-cell phenotyping based on marker combinations is a critical but challenging task due to (i) low reproducibility across experiments with manual thresholding, and, (ii) labor-intensive ground-truth expert annotation required for learning-based methods.

**Results:** We developed Tribus, an interactive knowledge-based classifier for multiplexed images and proteomic datasets that avoids hard-set thresholds and manual labeling. We demonstrated that Tribus recovers fine-grained cell types, matching the gold standard annotations by human experts. Additionally, Tribus can target ambiguous populations and discover phenotypically distinct cell subtypes. Through benchmarking against three similar methods in four public datasets with ground truth labels, we show that Tribus outperforms other methods in accuracy and computational efficiency, reducing runtime by an order of magnitude. Finally, we demonstrate the performance of Tribus in rapid and precise cell phenotyping with two large in-house whole-slide imaging datasets.

**Availability:** Tribus is available at https://github.com/farkkilab/tribus as an open-source Python package.

## Introduction

Multiplexed imaging techniques at single-cell resolution, such as tissue-based cyclic immunofluorescence (t-CyCIF) (Lin et al., 2018), co-detection by indexing (CODEX) (Black et al., 2021), and multiplexed ion beam imaging by time of flight (MIBI-TOF) (Keren et al., 2019), offer significant advantages for studying tissue architecture. These techniques enable researchers to measure dozens of proteins at single-cell resolution while preserving spatial origin information in tissue sections, providing novel insights into cellular phenotypes and tissue behaviors (Spitzer & Nolan, 2016). Multiplexed images require a sequence of processes to extract single-cell measurements, including image registration, stitching, cell segmentation, and quantification (Schapiro et al., 2022). Cell phenotyping is typically the final step before downstream analyses and often serves as the bottleneck in realizing the full potential of multiplexed images. The main challenges in cell-type phenotyping from multiplexed images include reproducibility limitations and unexpected marker combinations.

Classic methods for cell phenotyping such as manual gating and clustering, are reproducibility-limited. Gating (Staats et al., 2019) requires visualizing and manually setting marker expression thresholds for each marker in each sample. These hard thresholds are experiment-specific and can’t be reused across different batches. As the marker panel sizes and sample numbers increase, gating becomes time-consuming and unfeasible (Verschoor et al., 2015). Clustering algorithms, such as PhenoGraph (Levine et al., 2015), Leiden (Traag et al., 2019), and DBscan (Ester et al., 1996), have been applied for automatic data exploration (Liu et al., 2019). However, manual verification is still necessary to assign meaningful cell types to the resulting clusters. To achieve deeper profiling of cell types, over-clustering and subsequent merging of clusters is often required, a process that is both computationally intensive and time-consuming.

Several automated cell-type annotation approaches have been developed to overcome the reproducibility limits. For example, CellSighter is a supervised deep convolutional neural network-based algorithm for automatic cell phenotyping which requires expert-labeled images for training (Amitay et al., 2023). Another similar solution, MAPS is a supervised deep learning-based method that is computationally lighter than CellSighter (Shaban et al., 2024). These methods require pre-training on manually labeled datasets; however, the measured protein combinations (marker panel design) are experiment-specific, making it difficult to generate general reference populations for each cell type. In cases of unexpected marker combinations, the algorithms are required to be retrained in different scenarios.

To address the above challenges, we introduce a novel cell-type caller named Tribus, which incorporates the widely used self-organizing map (SOM) (Kohonen, 1982) unsupervised clustering method with a unique scoring function to assign cell types according to prior biological knowledge. Tribus requires only a cell measurement matrix and a prior knowledge table as inputs, without the need for training on expert annotations, and enables reproducible, automatic cell phenotyping across various multiplexed imaging and proteomic datasets. Tribus enables users to easily conduct analyses, visualize results, and perform quality control through an integrated Napari widget. We validate Tribus’s accuracy on four public multiplexed imaging and suspension mass cytometry datasets. We then compare Tribus’s performance to three other similar prior knowledge-based cell-type identification approaches: ACDC (Lee et al., 2017), Astir (Geuenich et al., 2021), and Scyan (Blampey et al., 2023), and demonstrate its utility in analyzing two large in-house t-CyCIF datasets. Tribus represents a novel user-friendly framework for semi-automated cell-type calling in multiplexed imaging and proteomics datasets.

## Methods

### Overview of Tribus

Tribus is a hierarchical framework for cell-type assignment in multiplexed image datasets based on prior panel knowledge (Fig 1). Running Tribus requires a marker expression table and a prior knowledge-based logic table. The logic table is defined as a data matrix *L* (*c*_*i*_, *m*_*j*_) containing values [−1,0,1], where *c*_*i*_ refers to the cell type *i*_*th*_ and *m*_*j*_ refers to the marker *j*_*th*_. If a marker is present in a specific cell type, the corresponding value in the logic table is assigned 1. If a marker is supposed to be absent in a specific cell type, the value in the logic table is set to −1. A score of 0 is assigned for neutral or unknown markers. Each cell type must have at least one positive marker in the logic table (Supplementary Table 1-3).

**Figure 1.**
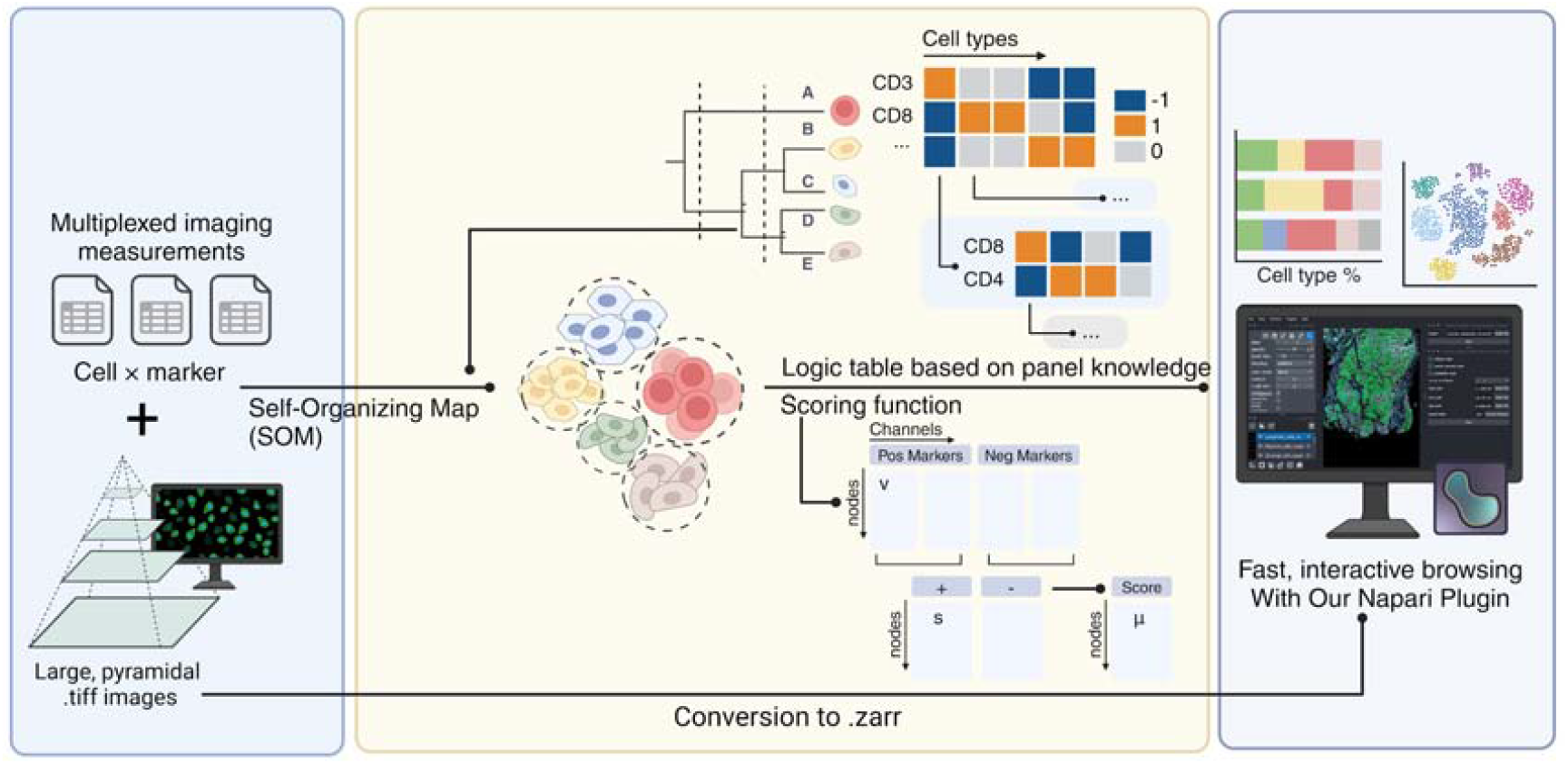
Overview of Tribus architecture. Tribus processes multiplexed imaging data by using a Self-Organizing Map (SOM) to cluster cells and a logic table based on panel knowledge to score and classify cell phenotypes. Results can be visualized with a Napari plugin for interactive exploration and quality control.

### Unsupervised clustering in Tribus

Tribus uses an unsupervised self-organizing map (SOM) method for clustering based on the MiniSOM package (Vettigli, 2013/2023). SOM can represent a high-dimensional input space as a map consisting of components called “nodes”. Quantization error (*Q*) was used to evaluate the algorithm performance, calculated by determining the average distance of the sample vectors 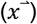 to the cluster centroids 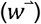.

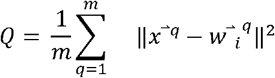

Quantization errors can only be compared under the same grid size (Pölzlbauer, n.d.). To determine the optimal grid size, we used the approach of Vesanto (Vesanto & Alhoniemi, 2000): 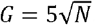, where *N* is the input data size (i.e., number of cells). The parameters of the SOM include *σ* (the spread of the neighborhood function) and the learning rate. Those parameters can be set by users or optimized by minimizing the quantization error with the hyperparameter tuning module based on the package hyperopt (Bergstra et al., 2013), where the objective function was to minimize the quantization error, and the search space for *σ* and the learning rate ranged from 0.001 to 5.

During analysis, Tribus first generates clusters/grids from the input data and then assigns each cluster to a certain cell type based on the logic table. Cell type assignment is performed hierarchically, meaning Tribus first assigns lower-level cell types followed by higher-level cell subtypes to create more precise categories. If the number of cells in the subset exceeds the user-defined threshold, Tribus will still generate clusters. Otherwise, Tribus directly calculates the scoring function for each cell type at the single-cell level.

### Cell type assignment by scoring functions

After SOM clustering, a node matrix *N* (*n*_*k*_,*v*_*j*_) is generated, where *v*_*j*_ is calculated as the median expression of the marker *m*_*j*_ in *n*_*k*_. We designed a scoring function based on the squared error concept, similar to the QueryStarPlot function of the FlowSOM package (Van Gassen et al., 2015). This function calculates the score *s*_*i*_ of a certain cell type *i* for each node *k*.

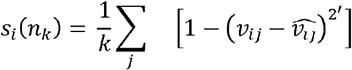

where,

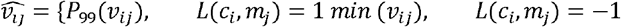

Instead of using the maximum value, the 99 percentile was chosen to make Tribus more robust to outliers. Assigning node *n*_*k*_ to cell type *c*_*i*_ using: 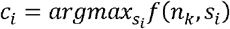. Note that the cell-type assignment process in Tribus accounts for ambiguous results. If the maximum score of a cluster falls below a certain threshold, the cluster is labeled as the “other” cell type. Similarly, if the difference between the maximum and second maximum score is smaller than a certain threshold, the cluster will be labeled as “undefined”.

### Napari plug-in

To efficiently evaluate cell-type labeling results, we developed a custom plugin integrated with the Napari (Ahlers, Jannis et al., 2023) framework. This plugin enables users to run Tribus on one sample at a time, display results simultaneously, or load previously saved data. We incorporated the ZARR format to overcome computational limitations associated with large datasets.

The key functionalities of the Napari plugin include:

i. Cell-type mask visualization: The plugin sorts and displays different cell-type labels as separate layers using visually distinct colors, allowing the user to overlay them with imaging data for quality control. This function is also available in a stand-alone Jupyter Notebook.
ii. Probability score visualization: cell masks are represented as a color gradient of the probability score assigned by the algorithm. This allows the user to identify and review ambiguous cells and assess the assigned “other” and “undefined” thresholds.
iii. Marker intensity visualization: The median expression levels of the selected markers are represented through gradient shading on the segmentation mask, enabling users to visually assess the results and identify potential biases.

### Methods for comparison and evaluation metrics

We compared the performance of Tribus with three similar prior knowledge-based cell-type calling tools: ACDC, Astir, and Scyan. We evaluated overall cell type annotation performance by comparing the Rand Index (Rand, 1971), accuracy, weighted F1 score (Powers, 2020), and Cohen’s kappa coefficient. We also compared the Matthews correlation coefficient (MCC), given the size variability of some cell types. All the above metrics were calculated using functions provided by Scikit-learn (Pedregosa et al., 2011).

### Benchmarking Datasets

#### Public datasets

We chose four public suspension mass cytometry and multiplexed imaging datasets with ground truth labels to test the performance of Tribus (Table 1). The AML dataset contains single-cell proteomic profiles of human bone marrow from patients with acute myeloid leukemia (AML) and healthy adult donors (Levine et al., 2015). The “NotDebrisSinglets” cell type was excluded from the analysis. The BMMC dataset was derived from bone marrow mononuclear cells (BMMCs) (Bendall et al., 2011). According to the research, Erythroblasts, megakaryocyte platelets, and myelocytes were merged as an unknown population and removed from the analysis. All “NotGated” cells were excluded (N = 61725 cells). The ductal carcinoma in situ (DCIS) dataset containing 79 clinically annotated surgical resections (Risom et al., 2022), including normal breast tissue (N = 9, reduction mammoplasty), primary DCIS (N = 58), and invasive breast cancer (IBC) (N = 12). Cell types were merged into endothelial, epithelial, fibroblast, immune, and myoepithelial during low-plex cell phenotyping. HuBMAP is a published co-detection by indexing (CODEX) imaging dataset (Hickey et al., 2023). Only donor 004 was manually annotated and, therefore, used in this study. For low-plex cell phenotyping, cell types were merged into epithelial, stromal, lymphoid, and myeloid.

#### In-house Datasets

We generated two in-house high-grade serous ovarian cancer (HGSC) datasets using t-CyCIF. The NACT dataset contains three images: two tumor sections after neoadjuvant chemotherapy (NACT) and one treatment-naive biopsy. The NACT dataset was stained using a 36-plex antibody panel (Supplementary Table 4). The Oncosys-Ova dataset contains 21 HGSC samples stained with a 14-plex panel (Supplementary Table 5). Marker expression tables were preprocessed using log transformation, z-score normalization, 99.9 percentile outlier removal, and co-factor 5 arcsinh transformation (Hickey et al., 2021) before phenotype assignment. In the NACT dataset, 976,082 cells were annotated using a cell phenotyping logic table based on the panel design. For the Oncosys-Ova dataset, approximately 10.5M cells were annotated. Following the ethical standards from the 1975 Declaration of Helsinki, every patient from ONCOSYS-Ova trials provided informed written consent to the collection, storage, and analysis of the samples and subsequent data. For the NACT dataset, the Mass General Brigham Institutional Review Board approved using human tissue samples. Informed consent was waived due to the use of archival samples and anonymization of the material.

## Results

### Tribus recovers fine-grained cell types as accurately as human experts

To evaluate Tribus’s ability to recover canonical cellular populations, we applied it to the benchmark DCIS dataset. Tribus successfully identified all populations highlighted in the study (Supplementary Fig. 1A), and the mean marker intensities of cellular populations were similar to those of human-labeled populations (Fig. 2A). One notable difference was observed in the MACS (macrophages) population, where Tribus labels displayed a lower median intensity of CD14 compared to manual gating. This discrepancy may be due to the fact that CD14 was not used for MACS identification in DCIS research, and thus CD14 was not constrained to be positive for MACS cells in the logic table. We mapped cell masks back to the original tiff images with a nine-color overlay of cell identity-related markers. The cell masks were consistent across all major cell types in Tribus labels and manual labels. False positive MACS in manual labels were corrected by Tribus (Fig 2B). When comparing manual labels and Tribus labels, UMAP visualization of all cell types from DCIS datasets showed that Tribus accurately annotated most cells (Fig. 2C). Using the manual labels as ground truth, Tribus achieved precision scores between 0.7 and 0.8, average recalls around 0.6, and F1 scores between 0.6 and 0.7 across most cell types (Supplementary Fig. 1B). These results suggest that Tribus can recover and annotate fine-grained cell types with a level of accuracy comparable to human experts.

**Figure 2.**
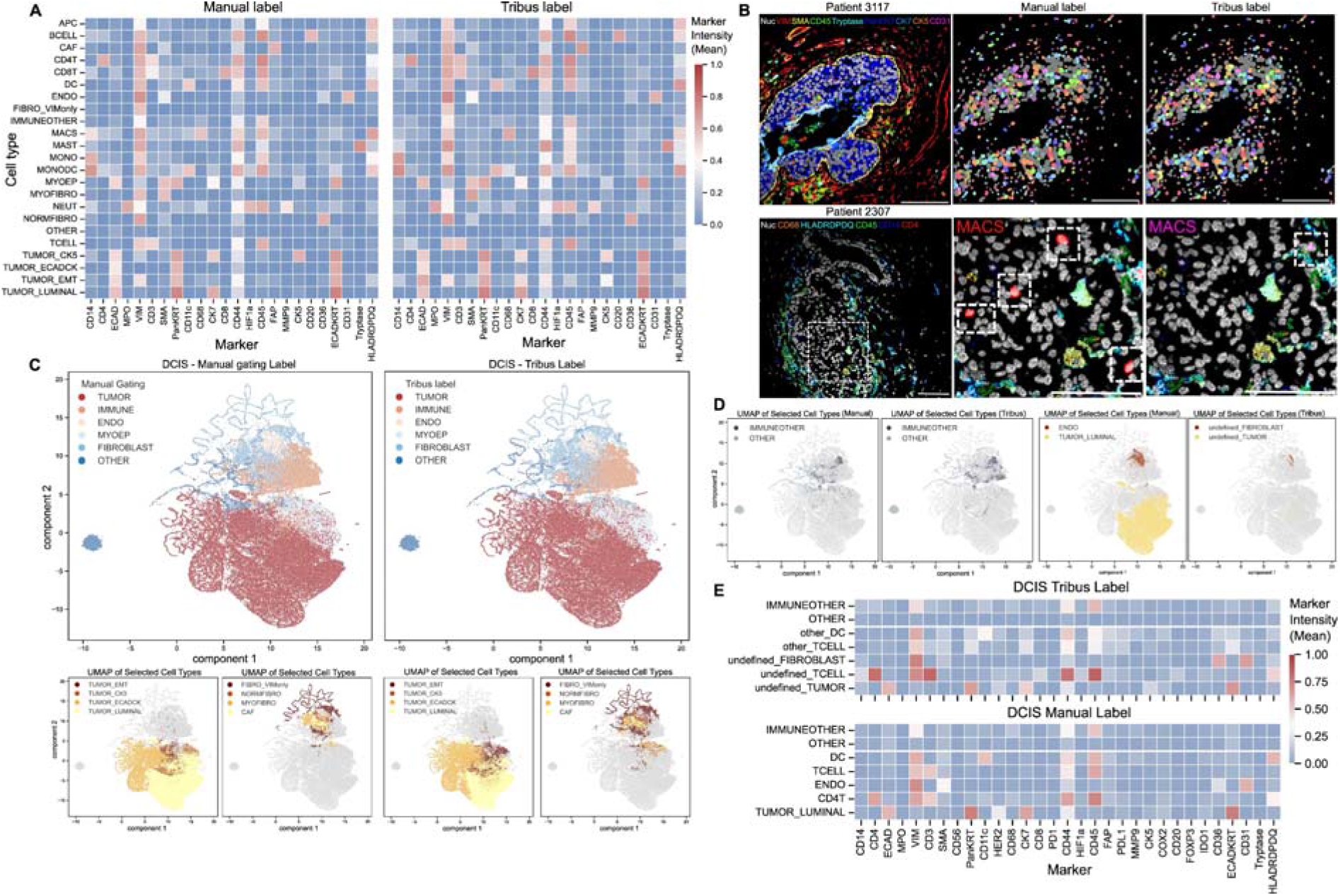
Tribus applied on the public DCIS dataset. (A) Heatmaps showing the mean marker intensity of manually gated cell populations and Tribus classification from the DCIS dataset are similar. (B) Representative MIBI images. The upper image is from patient 3117 (DCIS tumor) with a nine-color overlay of markers related to major cell types. Cell-type masks from the manual label and Tribus show few differences. The lower image is from patient 2307 (Normal tissue) with a six-color overlay of MACS-related markers. Compared to the MACS cell masks of manual and Tribus, some MACS cells from manual labels do not have related marker expression, which was corrected in Tribus annotation (canonical marker combinations were observed). (C) UMAP representation of manual and Tribus labels on the DCIS dataset. Cell types were color-coded based on original and Tribus annotation, the same cell type label was assigned the same color. (D) Comparing (1) undefined and other cell populations with ground truth labels, (2) undefined cellular subpopulations from Tribus annotation with relevant manual labeled cell populations. (E) Heatmaps comparing the mean marker intensity of manually gated cell populations and the ambiguous cell populations from Tribus annotation.

### Tribus can target ambiguous populations and discover phenotypically new subtypes

One challenge for prior knowledge-based cell-type calling methods is discovering ambiguous categories that are difficult to predefine in the logic table. For example, a group of unknown cells with low intensity across all markers. The DCIS dataset provides a good example for exploring ambiguous populations, as it includes the “IMMUNEOTHER” and “OTHER” cell types in the manual gating labels. The “OTHER” cell type has low intensities across all markers, while “IMMUNEOTHER” lacks specific immune subtype marker expression.

We verified that Tribus can successfully target ambiguous populations by adjusting the decision thresholds in the scoring function. We explored threshold settings and found Tribus is robust to different threshold values within a certain range (Supplementary Fig. 2A-B). We selected an undefined_threshold of 0.001 and an other_threshold of 0.04 for the analysis of the DCIS dataset. From the UMAP visualization, we observed that the same “IMMUNEOTHER” and “OTHER” clusters retained the same local structures (Fig. 2D). The marker expression heatmap demonstrates that Tribus-targeted ambiguous cell types share the same marker expression profiles as manually gated cell types (Fig. 2E). We also discovered new clusters, including undefined tumors, fibroblasts, T-cells, and DCs (Fig. 2D, Supplementary Fig. 1C). We found phenotypically new clusters of DCs, T-cells, and CD4 T cells using the mean marker expression heatmap (Fig. 2E). We observed novel marker expression combinations and cellular subtypes were validated by mapping cell masks back to the original images. We successfully identified CD36+CD31+ fibroblasts, HER2-luminal subtypes, and undefined T-cells which exhibit higher phenotypic marker expression than typical CD4 T-cells (Supplementary Fig. 3 A-C). These findings demonstrate that Tribus is not only effective in identifying ambiguous cell populations but also capable of discovering phenotypically novel cell subtypes. We suggest that Tribus could serve as a starting point for uncovering novel cell states.

### Tribus outperforms other similar methods

We benchmarked the performance of Tribus against three other approaches: ACDC, Scyan, and Astir. We chose these tools for benchmarking because they are all prior knowledge-based cell phenotyping methods, each designed for specific high-dimensional data types.

For AML and BMMC datasets, we used the knowledge tables provided by ACDC and Scyan. The knowledge table was reformatted into the logic tables/YAML files accordingly for Tribus/Astir. Since no knowledge tables were available for the DCIS and HubMAP datasets, we generated them for all methods based on the panel information provided in the original study. The ACDC analysis was performed with the parameters (n_neighbor = 10, thres = 0.5) from the example scripts. The parameters for the Astir analysis were chosen (max_epochs = 1000, learning_rate = 2e-3, initial_epochs = 3) based on the tutorial provided in Colab. The parameters for Scyan analysis were chosen based on the example provided in the GitHub repository.

Benchmarking experiments showed that Tribus outperformed the other methods in terms of efficiency and accuracy. Tribus outperforms Astir, ACDC, and Scyan methods across metrics on highly multiplexed imaging datasets, DCIS and HubMAP, in both high- and low-plex cell phenotyping. Tribus has comparable performance on suspension mass cytometry AML and BMMC datasets compared to ACDC and Scyan, which were designed specifically for mass cytometry datasets (Fig. 3A). On all benchmarking datasets, Tribus exhibits a running time shorter by an order of magnitude compared to other methods (Fig. 3B). Tribus successfully identified all cell types highlighted in the four public datasets (Supplementary Fig. 4 A-D).

**Figure 3.**
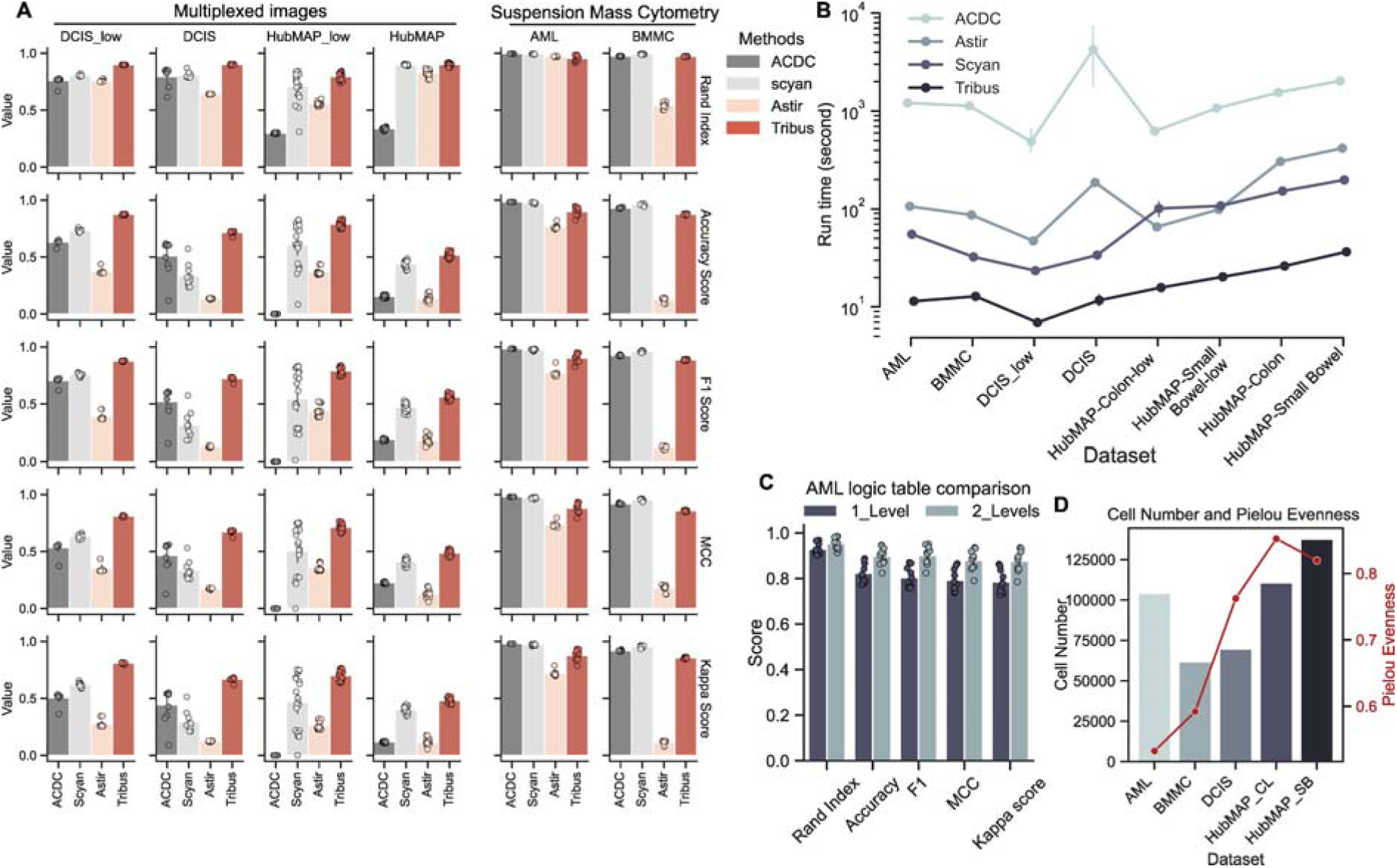
(A) Performance comparison of Tribus and other similar methods on four datasets (AML, BMMC, DCIS-lowplex, DCIS, HubMAP-lowplex, HubMAP) using five metrics for each. All analyses were repeated ten times. Using standard deviation for the error bar. (B) Models running time comparison. All analyses were repeated ten times and used standard deviation as the error bar. (C) Compare Tribus performance under different logic tables for the AML dataset, using standard deviation as the error bar. Each analysis was repeated ten times. (D) Data complexity comparison over public benchmarking datasets, showing the number of cells and Pielou’s evenness index. Higher Pielou’s evenness index represents high diversity and high evenness of cell populations.

We then explored how the structure of the logic table influenced performance. Using the AML dataset as an example, we applied Tribus analysis with (1) a logic table with only one global level and (2) a logic table that includes major cell types at the global level and sub-phenotypes (for example, CD16- and CD16+ NK cells) at the second level (Supplementary Table 1-3). We repeated the Tribus analysis ten times, calculated the average metrics, and visualized the results for each logic table configuration. When adjusting the logic table for the AML dataset, using a hierarchical logic table improved accuracy and increased the F1 score by 0.1 (Fig. 3C).

Finally, we calculated Pielou’s evenness index to illustrate the increasing complexity across benchmarking datasets (Fig. 3D). Tribus’s performance remained relatively good and stable from suspended single-cell bone marrow datasets to highly complex human intestine slide datasets. In summary, Tribus outperformed other similar methods in both accuracy and efficiency, particularly in highly multiplexed imaging datasets, while maintaining robust results across different cell phenotyping contexts.

### Tribus yields rapid and accurate cell phenotyping in large whole-slide image datasets

We evaluated the performance of Tribus on large in-house multiplexed image datasets. For the Oncosys-Ova dataset, we used a one-level logic table consistently across all 21 samples (Supplementary Fig. 5A). We applied a four-level logic table to the NACT dataset (Supplementary Fig. 5B).

In the NACT dataset, Tribus successfully identified major cell phenotypes and subtypes, which we characterized based on the available marker panel (Fig. 4A). The UMAP projection showed distinct cell-type populations in the NACT dataset (Fig. 4B-C) and minimal batch effects in phenotyping analysis (Supplementary Fig. 5C). The multiplexed t-CyCIF image from sample 06 of the NACT dataset displayed nuclei and representative tumor and immune markers. Tribus accurately identified the CD20+, CD8a+, and CD4+ cell populations within a Tertiary Lymphoid Structure (TLS) despite the dense organization of these sub-phenotypes (Fig. 4D). Tribus also annotated the proliferating subpopulation of tumor cells and tumor-infiltrating CD8+ cells from a dense area (Supplementary Fig. 5D), indicating that Tribus can generate accurate phenotype labels in complex tissue architectures.

**Figure 4.**
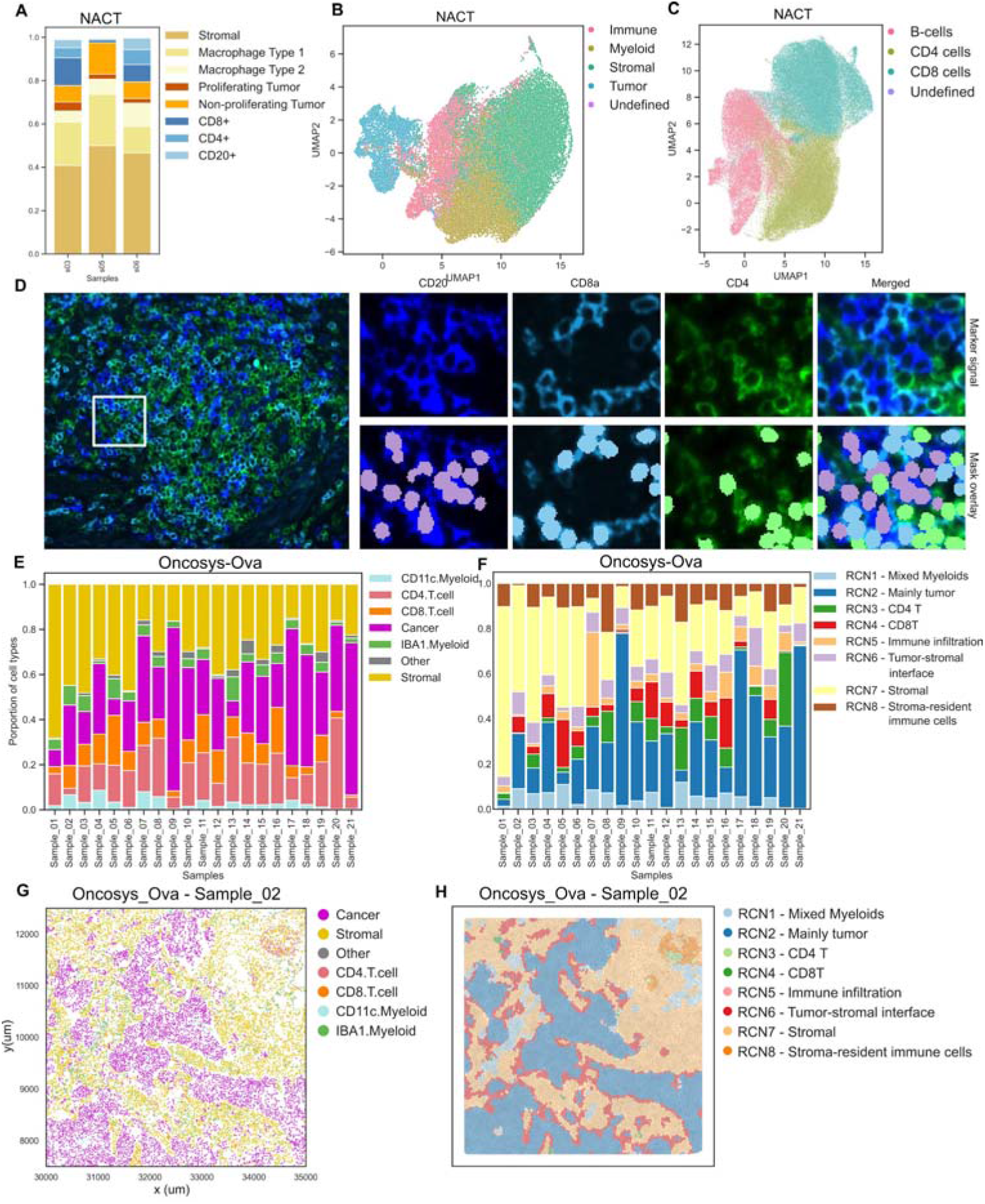
(A) Annotated barplot of the cell phenotype compositions per sample in the NACT dataset. (B) UMAP visualizes distinct cell populations colored by the major cell types in the NACT dataset. (C) UMAP visualizes distinct cell populations colored by the immune subtypes in the NACT dataset. (D) Representative tCyCIF image (sample 06) of the NACT dataset, showing the nuclei and representative tumor and immune markers. Tribus accurately identified the CD20+, CD8a+, and CD4+ populations within a Tertiary Lymphoid Structure (TLS) despite the dense organization of these sub-phenotypes. (E) The stacked barplot shows cellular proportion in all samples. (F) Stacked barplot shows various RCN proportions in samples. (G) The scatter plots show the local tissue architecture colored by cell types. (H) The Voronoi plot visualizes the structures of RCNs in the corresponding region of Figure G.

In the Oncosys-Ova dataset, Tribus accurately identified six cell types with substantial cellular proportion heterogeneity among samples (Fig. 4E). UMAP projections showed separated cell populations and low batch effects (Supplementary Fig. 6A). The heatmap showed canonical marker expression combinations for each identified cellular population (Supplementary Fig. 6B). We used Scimap (Nirmal & Sorger, 2024) to calculate the fractions of neighboring cell types within a radius of 100 μm, then applied k-means clustering (k=10) on the neighborhood matrix and generated eight Recurrent cellular neighborhoods (RCN) (Fig. 4F). The RCNs are distinct spatial domains within the tissue and successfully capture relevant spatial biology based on Tribus cell phenotypes (Supplementary Fig. 6C). The representative presentation of the RCNs across tissue uncovered the complex tissue architecture such as the tumor-stromal interface and tumor-infiltrating lymphocytes cells (Supplementary Fig. 6D). Tribus-based spatial analysis enabled us to map the tumor-stromal interface in complex tumor-rich regions (Fig. 4G, Supplementary Fig.6D) and plot the stroma-resident immune cells in a stromal-rich region (Supplementary Fig.6D). These results suggested that Tribus adapts well in the workflow of cell phenotyping on large whole-slide images and downstream spatial pattern analysis.

## DISCUSSION

Cell-type calling is a crucial step in high-dimensional image analysis. The growing complexity and increasing number of panels in high-dimensional data necessitate the development of reproducible and automated cell-type calling approaches. In this study, we developed Tribus, a semi-supervised cell-type calling analysis framework for multiplexed imaging datasets. Tribus offers advantages in efficiency, accuracy, user-friendliness, and reproducibility without the need for training using manual labels.

Tribus was generated as an automatic cell-type caller for high-dimensional multiplexed imaging data, incorporating biological knowledge from the panel design into the analysis. This was achieved through carefully designed scoring functions based on marker expression per grid, generated by unsupervised clustering to minimize bias. When the number of cells in the clusters was below the user-defined threshold, Tribus skipped generating the clusters and calculated scores at the single-cell level. This flexible scoring function calculation strategy enabled the discovery of rare cell types. Tribus was integrated with Napari, and a plugin was provided to enable one-click import of the cell-type identification results, significantly enhancing the simplicity of interactive quality control. This integration allows for convenient operation by users who are unfamiliar with programming.

Cell-type annotation from bioimages presents inherent challenges due to imperfect cell segmentation and collateral spillover. Expanding nuclei masks by a few pixels can enhance cytoplasmic marker visibility, as it allows better signal capture when cells express these markers. However, in dense tissue areas, this might increase spillover, highlighting the importance of the hierarchically structured logic table. Typically, mild spillover affects only part of a cell, whereas a true signal produces a more uniform expression pattern and a higher mean fluorescence intensity. Tribus was designed to account for both positive and negative components of the expected marker expression, and it also includes the option to set markers with expected false-positive expressions as neutral. Thus, careful design of the logic table and its hierarchy in Tribus can aid in cell phenotypic separation, as only a subset of cells is considered at the lower hierarchy levels during phenotype assignment.

However, Tribus is not without limitations. The performance of Tribus is strongly tied to the quality of the input dataset and the prior knowledge of expected cell types in the user-defined initial logic table. To assign a uniform logic table, the samples should have even staining patterns both within and across slides. Uneven staining patterns and antibodies with suboptimal signal-to-noise ratios can significantly affect the results. For such suboptimal datasets, users can create a hierarchical logic table where major cell phenotypes with clear marker signals are separated at higher levels. This can result in more accurate labeling if distinct areas of the image are affected. Additionally, Tribus cannot identify cell types not included in the input logic table, but it can return undefined-or other-cell types for further exploration.

Overall, we propose Tribus as a fast, accurate, and user-friendly cell-type identification method that can be integrated into multiplexed image analysis frameworks.

## Author contributions statement

Z.K., A.S., T.F., and J.C. conceptualized, designed, and implemented the algorithm and software. Z.K. performed benchmarking analyses and analyzed the Oncosys-Ova dataset. A.S. analyzed the NACT in-house dataset. Z.K. and A.S. wrote the manuscript. I-M.L, F.P., E.A., and K.E. generated the in-house HRP dataset. A.V., U-M.H collected Oncosys-Ova samples. A.J., S.S. stained the Oncosys-Ova samples and performed the t-CyCIF experiment. A.F. conceived and supervised the study, provided resources, and wrote the manuscript. All authors have reviewed and approved the manuscript.

## Acknowledgments

We thank Prof. Sampsa Hautaniemi for his helpful feedback on the manuscript and the IT Center for Science (CSC) for the computational resources. The graphical overview was created with Biorender.com.

## Data and code availability

The Tribus package is publicly available on Github: https://github.com/farkkilab/tribus. The datasets and scripts used to repeat the results are publicly available at Synapse: https://www.synapse.org/Synapse:syn53754523/files/.

